# A framework for predicting soft-fruit yields and phenology using embedded, networked microsensors, coupled weather models and machine-learning techniques

**DOI:** 10.1101/565010

**Authors:** Mark A. Lee, Angelo Monteiro, Andrew Barclay, Jon Marcar, Mirena Miteva-Neagu, Joe Parker

## Abstract

Predicting harvest timing is a key challenge to sustainably develop soft fruit farming and reduce food waste. Soft fruits are perishable, high-value and seasonal, and sales prices are typically time-sensitive. In addition, fruit harvesting is labour-intensive and increasingly expensive making accurate phenological predictions valuable for growers. A novel approach for predicting soft fruit phenology and yields was developed and tested, using strawberries as the model crop. Seedlings were planted in polytunnels, and environmental and yield data were collected throughout the growing season. Over 1.2 million datapoints were collected by networked microsensors which measured spatial and temporal variability in air temperature, relative humidity (RH), soil moisture and photosynthetically active radiation (PAR). Fleeces were added to a subset of the plants to generate additional within-polytunnel variation. Cumulative fruit yields followed logistic growth curves and the coefficients of these curves were dependent on micro-climatic growing conditions. After 10,000 iterations, machine learning revealed that RH was the optimal factor informing the coefficients of these curves, perhaps because it is an integrative metric of air temperature and water status. Trigonometric models transformed weather forecasts, which showed a relatively low agreement with polytunnel air temperature (R^2^ = 0.6) and RH (R^2^ = 0.5) measurements, into more accurate polytunnel-specific predictions for temperature and RH (both R^2^ = 0.8). We present a framework for using machine-learning techniques to calculate curve coefficients and parametrise coupled weather models which can predict fruit yields and timing to a greater degree of accuracy that previously possible. Dataloggers measuring environmental and yield data could infer model parameters using iterative training for novel fruit varieties or crop types growing in different locations without *a-priori* phenological information. At this stage in the development of artificial intelligence and networked microsensors, this is a step forward in generating bespoke phenological prediction models to inform and support growers.

## 1. Introduction

Soft fruits are economically valuable and are key contributors to global food security, particularly because they are important sources of micronutrients, antioxidants and fibre (Tilman and Clark 2014). For example, in 2016, 9.1 million tonnes of commercial strawberries (*Fragaria × ananassa* Duch.) were grown worldwide with a value of approximately $18 billion. Countries across Asia (4.7 million tonnes), the Americas (2.1 million tonnes), Europe (1.7 million tonnes) and Africa (0.6 million tonnes) all dedicated substantial land areas to production (FAOSTAT 2018). China, USA, Mexico, Egypt, Turkey, Spain, Russia, Poland, Republic of Korea, Japan, Germany, Morocco, Italy and the UK each produced more than 100,000 tonnes of strawberries in 2016, with China and USA each producing approximately 3.8 million tonnes (FAOSTAT 2018).

Predicting fruit yields, fruit quality and plant phenology is a key agronomic challenge for many soft-fruit growers, since harvesting these crops is labour-intensive and based on increasingly expensive ‘just-in-time’ staffing. Soft fruit crops are perishable, high-value and seasonal, and sales prices are typically agreed in advance and are time-sensitive, making accurate phenological predictions incredibly valuable for growers (Morris et al. 2017). Air temperatures have been correlated with strawberry phenology and fruit quality in previous work, and is the most commonly used parameter in current predictive phenological models (Sønsteby et al. 2017). Relatively basic and inaccurate models continue to be used by growers to predict the onset of life cycle stages (e.g. truss formation, flower induction, 25% fruit development and 50% fruit development). These models include growing degree-hours (GDH) and growing degree-days (GDD) whereby plant development is tracked based on the number of hours or days where the mean temperature is above a threshold value or within a given range of values (Tanino and Wang 2008; Krüger et al. 2012). GDH and GDD are used by some growers to plan staff recruitment, customer contracts and management interventions which can then be used to speed up or slow down ripening (e.g. venting, opening doors and applying fleeces). GDD and GDH models are imprecise, require an accurate *a-priori* value for the GDH or GDD of a given variety and predictions are highly dependent on speculating on future growing conditions using weather forecasts. These forecasts are unlikely to be locally accurate since the weather stations are often considerable distances from a farm and polytunnels generate their own microclimates.

Many soft fruits, including strawberries, are now grown under polytunnels or in glasshouses, with the warmer conditions allowing the growing seasons to be extended (Salamé-Donoso et al. 2010; Vicente et al. 2014). When growing conditions exceed threshold values (e.g. a maximum air temperature or minimum soil moisture content) then physiological stress can reduced fruit yields (Grant et al. 2016). Reductions in sunlight, abrupt changes in air temperature and freezing, as well as periods of drought have also been associated with changes in strawberry yields (Ariza et al. 2015; Martínez-Ferri et al. 2016), whilst adequate soil temperatures are a crucial aspect of germination (Domínguez et al. 2014). Genomic analysis of the strawberry revealed genes critical to valuable horticultural traits including flavour, nutritional value and flowering time, but the interactions of these genes with environmental cues remains poorly understood (Shulaev et al. 2011). Strawberry cultivars are highly variable and their responses to growing conditions will be cultivar specific (Sønsteby and Heide 2008). Predictive phenological models are therefore required to be cultivar specific and their accuracy are dependent on data availability and quality, as well as the modelling approach.

A new generation of miniaturised autonomous real-time environmental sensors and artificial intelligence offers the potential to link local microclimate conditions with mesoscale weather forecasts to more accurately model microclimates for each crop up to a fortnight in advance. To utilise this technology to optimise management for predictable yields and ripening schedules in soft fruits, a new modelling approach is proposed, and is tested here, which combines locally-fitted weather station data with logistic growth curve and delay functions to predict strawberry ripening. These sensors’ high spatiotemporal resolution offers the potential to build empirically parameterized models using machine learning to optimise crop management at fine spatial scales. Machine learning techniques have been previously applied to community composition analyses (Valle et al. 2014), environmental science (Jung et al. 2010) and artificial intelligence (Ghahramani 2015). Such techniques offer many benefits over more traditional modelling approaches; data may be highly dimensional, heterogeneous, and contain spatiotemporal structure, but are still able to generate useful location-specific models with minimal human input (Sukumaran et al. 2016). Owing to the rapidity with which this technology is developing, we are not aware of any industrial or academic studies which have applied machine learning to crop yield optimisation and predictions of plant phenology.

We investigated the efficacy of a new modelling framework for predictions of soft fruit phenology and yields, using strawberry plants as our model crop. We collected environmental and yield data throughout the growing season using high resolution networked microsensors (Figure 1a and 1b). We combined first principal models, such as known meteorological patterns (e.g. cyclical and seasonal variation in temperatures) and developed models simulating site-specific polytunnel microclimates. We also tested established biological models to predict fruiting (e.g. logistic growth curves). We aimed to improve our understanding of soft fruit phenology and provide a way forward in the use of ‘big data’ to parametrise machine learning tools to support the sustainable development of the soft fruit industry (Figure 1c).

**Figure 1:**
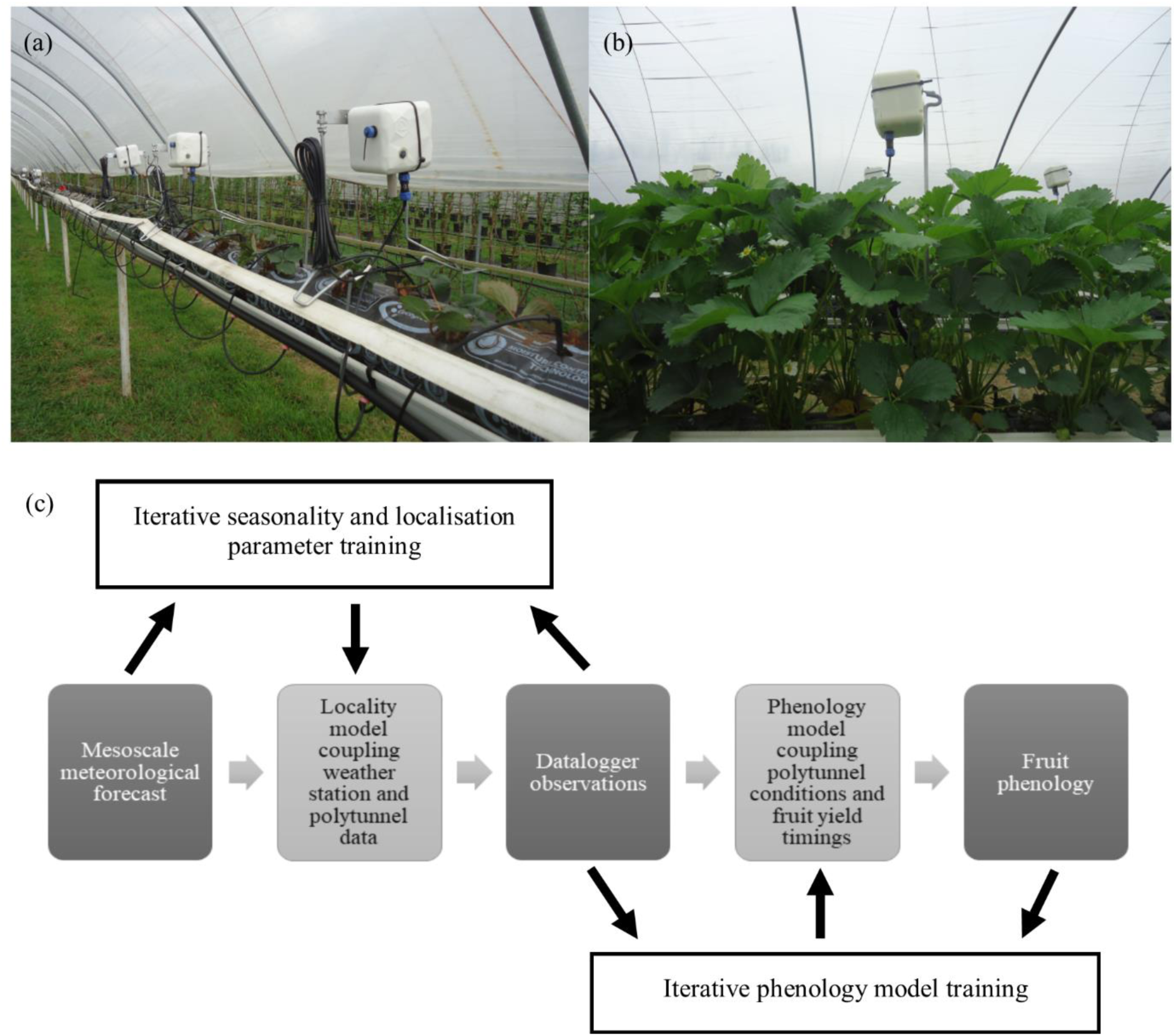
Photos of experimental setup showing (a) dataloggers (N = 18) in the polytunnel as strawberry seedlings are planted at the beginning of the study and (b) dataloggers raised above the plants during the experiment. Dataloggers measured air temperature, relative humidity, PAR irradiance and soil moisture and (c) these data were used to test a novel framework for using machine learning to predict fruit phenology.

We hypothesised that:

1. Cumulative fruiting mass follows a logistic growth curve over time.
2. The coefficients of these curves are dependent on growing conditions.
3. Growing conditions are predictable using trigonometric transformed weather forecasts.
4. Coefficients can be calculated using machine learning to predict phenology and yields.

## 2. Materials and Methods

### 2.1. Site

The experiment was conducted under a polytunnel (120m × 7m) at Berry Gardens™ research facility, Clock House Farm, Coxheath, UK (51.23N, 0.50E, elevation = 130m) between March and July 2017. The polytunnel was orientated North-South with surrounding polytunnels growing different strawberry varieties and other soft fruits. Strawberry plants, which had been cold stored as tray plants (Variety: “Malling Centenary”), were planted in 1.2m raised beds containing a coco peat substrate on 31^st^ March 2017. Plants were propagated in an off-site nursery, before being cold stored and delivered in a dormant state. Eight plants were randomly assigned and planted in 18 growbags of substrate (1m in length) so that there was a total of 144 plants in the experiment. Water availability was monitored and regulated by measuring the moisture content of the substrate using a handheld probe (Delta-T WET probe: Cambridge, UK) on a daily basis and the speed of water inflow from a centralised irrigation system was adjusted to achieve a constant level of water availability over time. The crop was subjected to a conventional pest and disease management strategy, integrating permitted agrochemicals and beneficial insects.

### 2.2. Automated environmental data collection

A wireless networked environmental sensor and datalogger system was installed prior to planting which comprised 18 integrated dataloggers (Mothive Ltd: Kingston, UK). One datalogger was assigned to each of the growbags (containing 8 plants) so that there was a total of 18 loggers with each collecting data using four environmental sensors (Figure 1a). Each experimental growbag was separated by another growbag of strawberry plants which were not part of the experiment. The system recorded four variables every eight minutes for each datalogger throughout the experiment and sent these data directly to the cloud for storage. During the experiment, over 1.2 million datapoints were collected across all the dataloggers. Datalogger were mounted on metal posts and raised above the plants (Figure 1b). During the experiment plant heights were monitored and the height of the loggers adjusted to prevent interference with the plants or shading of the leaves. Each datalogger had sensors which measured soil moisture, photosynthetically active radiation (PAR) irradiance, air temperature and relative humidity. The sensors were located close to the plants, and raised regularly to account for plant growth, so that the measurements were representative of growing conditions. Weather data was obtained from a weather station located approximately 3 miles NNW of the site (51.26N, 0.47E, elevation = 50m). The weather station recorded air temperature, relative humidity, wind speed, cloud cover and atmospheric pressure every two hours.

### 2.3. Experimental treatments

To investigate the effects of management interventions on growing conditions, plant development and phenology, semi-transparent cotton fleeces were applied to a subset of the strawberry plants from 26^th^ April until harvest. There were three fleece management treatments; (a) semi-permanent fleecing, whereby fleeces were only removed when the air temperature exceeded 30 °C and reapplied when the air temperature dropped below 30 °C (hereafter termed “permanent”), (b) managed fleecing, whereby fleeces were removed when the air temperature exceeded 24 °C and reapplied when the air temperature dropped below 24 °C (hereafter termed “managed”), and (c) control, whereby fleeces were never applied. Treatments were located more than 10m from the polytunnel door to avoid capturing edge effects and were separated by approximately 10m to avoid any interactions between treatments.

The networked sensor and datalogger system raised alerts and the fleeces were removed in the managed and semi-permanent treatments when the mean air temperature across the six control dataloggers exceeded 24 °C or 30 °C for five consecutive measurements. The fleece was then reapplied when the mean air temperature dropped below the thresholds for five consecutive measurements. This avoided excessive removal and reapplication of the fleeces which may have damaged the plants. The sensors were located below the fleece so that measurements were representative of the growing conditions. We also investigated the effect of row by separating our dataset into the central and leg rows within the polytunnel. This resulted in a factorial design with row and fleecing treatment as the two main categorical variables. Treatments also served to generate variation in fruit yields, phenology and quality to incorporate into the machine learning framework.

### 2.4. Strawberry yields, quality and plant physiology

Plants were monitored daily during the experimental period to establish when the fruits would be suitable for picking. Fruit picking commenced on 30^th^ May 2017, two months after planting. Picking then occurred every two days until the final fruits were picked on 11^th^ July 2017. During this period, measurements were taken from fruits picked from each of the eight plants that were selected for harvest in each growbag and hence for each of the 18 dataloggers. Fruit yield and quality was assessed by measurements of total fruit yield, total class 1 yield, mean mass of five randomly selected class 1 fruits and mean sugar content of five randomly selected fruits. Sugar content was measured using a handheld refractometer reading the percentage sugar content in degrees Brix (°Brix). Fruits were selected for picking based on their suitability for market as assessed by experienced fruit pickers using a standard commercial scale.

Physiological measurements of the plants occurred during three surveys over the experimental period on 18^th^ May, 26^th^ May and 9^th^ June. Stomatal conductance was measured from one randomly selected plant from each growbag using a SC-1 Leaf Porometer (Labcell Ltd: Alton, UK) and at that time soil temperature was also measured using a soil thermometer. Prior to the measurements taking place the porometer was left to equilibrate to polytunnel conditions for 30 minutes and calibrated. On each of the survey dates, stomatal conductance measurements were taken from plants in each growbag at 10:00 and then this was repeated until 13:00. This meant that there were between 3 and 6 measurements from different plants for each datalogger. In the case of the fleeced treatment, measurements took place under the fleece without removing it to quantify the effect of the fleece. If the fleeces were removed because temperatures exceeded the threshold values, then measurements continued. This assessed the direct effects of the fleeces on environmental conditions and stomatal conductance.

Fruit yields and quality as well as the physiological measurements for each growbag and sensor were time stamped so that these values could be compared with the air temperature, relative humidity, soil moisture and irradiance measurements as recorded by the environmental sensor system at that time. This also allowed for phenology to be assessed over the duration of the experiment.

### 2.5. Statistics and coupled growing conditions modelling

Generalised linear models (GLMs) were used the assess the effects of fleece treatments and rows on growing conditions within the polytunnel. These were included as categorical explanatory variables in statistical models as well as a binary variable describing whether the fleece was off or on, with analyses carried out separately for air temperature, relative humidity, PAR and soil moisture as response variables. Variation over time was also included in statistical models as two continuous explanatory variables (time of day and days since planting), with the shapes of the relationships being identified using linear, quadratic and trigonometric functions. The optimal models were then selected based on maximum r^2^ values. The experimental design resulted in multiple measurements being taken from the same location which necessitated inclusion of day and time within the statistical models, alongside the treatments. Measurements taken every eight minutes for each variable resulted in a large dataset, favouring low P values. Significant results (P < 0.05) were therefore assessed in relation to their relative effect size (see Results section).

Weather station data were compared with data recorded by the dataloggers from the control treatment for two key parameters, air temperature and RH. Since these parameters were measured every eight minutes by the environmental sensor system and every two hours by the weather station, data were averaged over two hours. For both parameters, linear regression was used to correlate polytunnel conditions with weather station conditions, with the degree of fit of the regression line again being assessed using r^2^. Residual differences between the weather station and polytunnel conditions (Residual = polytunnel value – weather station value) were then plotted over time. Linear, quadratic and trigonometric functions were used to describe these residuals, with the optimal function being chosen based on r^2^ values. The equation of the fitted linear regression between the weather station and polytunnel, and then the equation of the optimal residual relationships was used to transform weather station values into simulated polytunnel conditions. These simulations were then compared to measured polytunnel data using linear models, with the degree of fit being assessed using r^2^.

Fruit yields and quality were assessed using GLMs, with fleece treatments and rows included as fixed effects and harvest day included as a random effect. This accounted for multiple measurements being taken from the same location and avoided pseudo-replication. Each datapoint represented mean fruit yield per plant, mean mass per fruit or sugar content for one growbag and hence one of six dataloggers for each treatment and for each harvest. Values were therefore mean values of the eight plants assigned to each datalogger for each harvest day. The effects of fleece and row on stomatal conductance measurements were assessed using GLM with a gaussian error structure. LME was then used to quantify the effects of polytunnel environmental conditions on rates of stomatal conductance.

To assess the development of fruits over time, logistic growth curves were used the describe the accumulation of fruit over time by fitting logistic growth curves to cumulative fruit yields per plant and cumulative class 1 fruit yields per plant. These were fitted separately for each treatment since significant treatment effects were demonstrated using GLM. Logistic growth models were fitted to the dataset using the ‘fit_growthmodel’ function from the ‘growthrates’ package in R. Initial model parameters were manually added, and optimal coefficients subsequently generated, with the degree of fit being assessed using r^2^ values. In terms of fruit quality, variation in fruit sugar content was assessed using linear models. All statistical analyses and data modelling were carried out using the statistical software, R (R Core Team, v3.4.1).

### 2.6. Fruit phenology machine learning approach

The logistic function for cumulative fruit yields after time *t*, *y(t),* was modified by the addition of a delay parameter, *d*, representing the initial vegetative growth of the strawberry plants (equation 1). Where *r* = the intrinsic growth factor; *K* = the maximum possible yield (carrying capacity); and *p=2* (a tuning parameter). This function was numerically optimised using gradient descent with an untuned normally-distributed proposal mechanism (a_i+1_ ~ a_i_ + *N*(1,(a_i_*0.01) for *r* and *d*; a_i+1_ ~ a_i_ + *N*(1,(a_i_*0.001) for *K*). Optimisations were run separately for each time-series, for 500 and 10,000 steps, using 10 replicates to check convergence. Each datalogger represented an experimental replicate for the machine learning optimisation (N = 18) with microclimatic variation generated by natural variation within the polytunnel as well as variation derived from the fleece and row treatments.

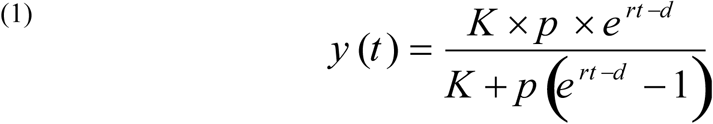

## 3. Results and discussion

Seedlings were planted on 31^st^ March 2017 and data was collected until the final fruit harvest on 11^th^ July 2017. Polytunnel air temperatures generally increased over the duration of the study, with the mean daily air temperature of 13.1 ± 0.05°C in April gradually rising to 22.4 ± 0.08°C in July. Air temperatures fluctuated cyclically during the day, with the maximum daily temperature most frequently falling between 14:00 and 15:00 and the minimum daily temperature falling between 04:00 and 05:00. Soil moisture remained relatively static over the period, with mean soil moisture in April, June and July being 55.0 ± 0.06%, 54.6 ± 0.04% and 49.7 ± 0.2%, respectively. However, in May, soil moisture declined for several days with mean daily values dropping below 40% on five days and below 45% on seven days resulting in a lower mean soil moisture content of 49.7 ± 0.08%. PAR irradiance followed a cyclical daily pattern, and peak values occurred most frequently between 14:00 and 15:00 with night time values of ~0 W m^-2^ recorded approximately between 21:00 and 05:00, depending on the day length. RH also followed a cyclical daily pattern but declined over time from a mean daily RH of 71.3 ± 0.1% in April to 64.5 ± 0.3% in July. Minimum values for RH occurred most frequently between 14:00 and 15:00 with maximum values occurring between 05:00 and 06:00, an inverse pattern to the timings of maximum and minimum air temperatures.

### 3.1. Parameterisation of polytunnel conditions

The models which optimally explained variation in air temperatures, RH, PAR irradiance all included the fleece treatments, the row of the polytunnel, linear parameters which described variation over the period (i.e. between days) and trigonometric parameters which described the cyclical nature of measurements taken throughout the day (Table 1). Trigonometric functions were associated with the greatest effect sizes and explained the most variation in the datasets for air temperatures, RH and PAR irradiance. In the case of soil moisture, however, a positive linear function was a superior parameter for explaining variation throughout the day, likely because this was manually adjusted daily.

**Table 1:**
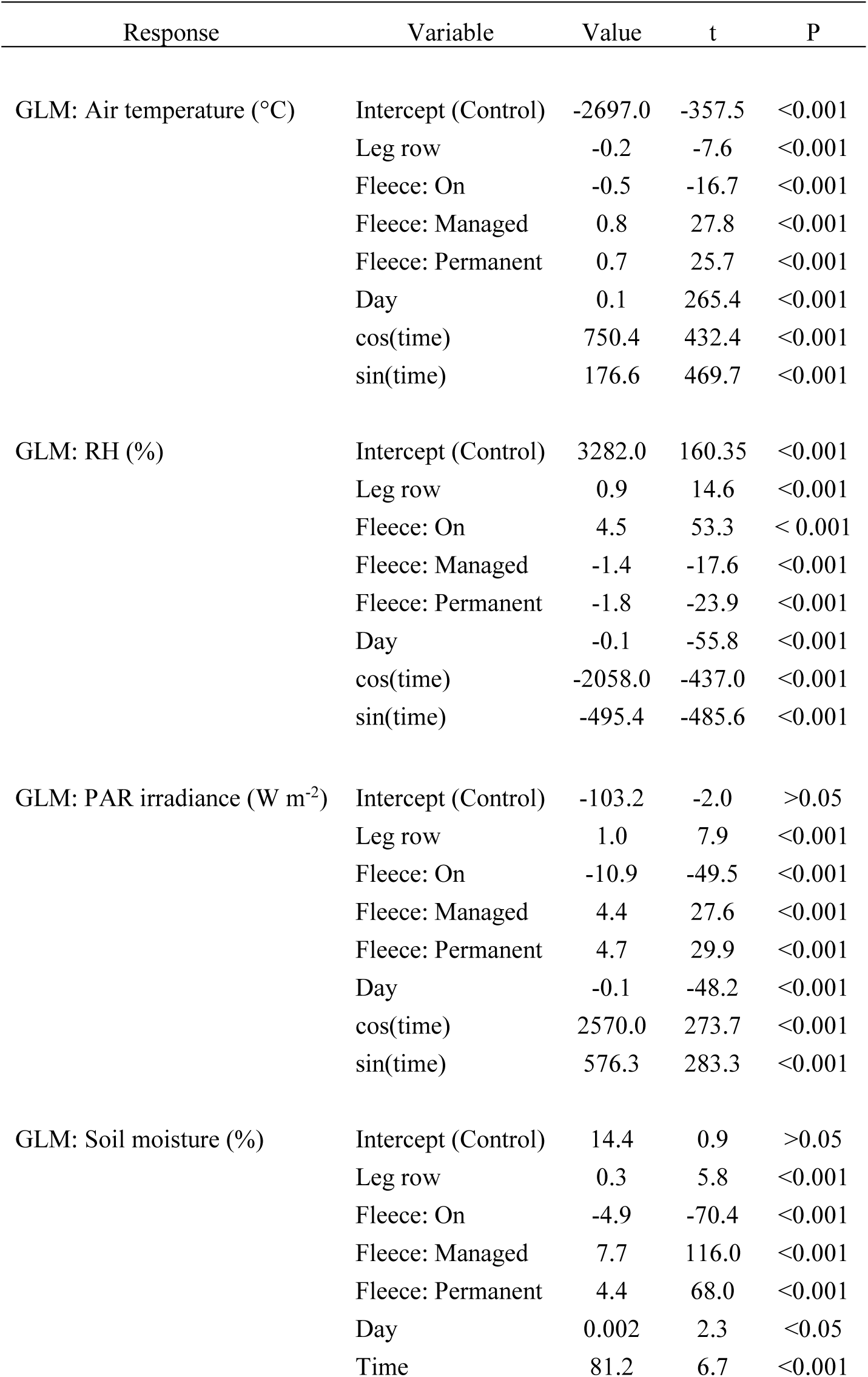
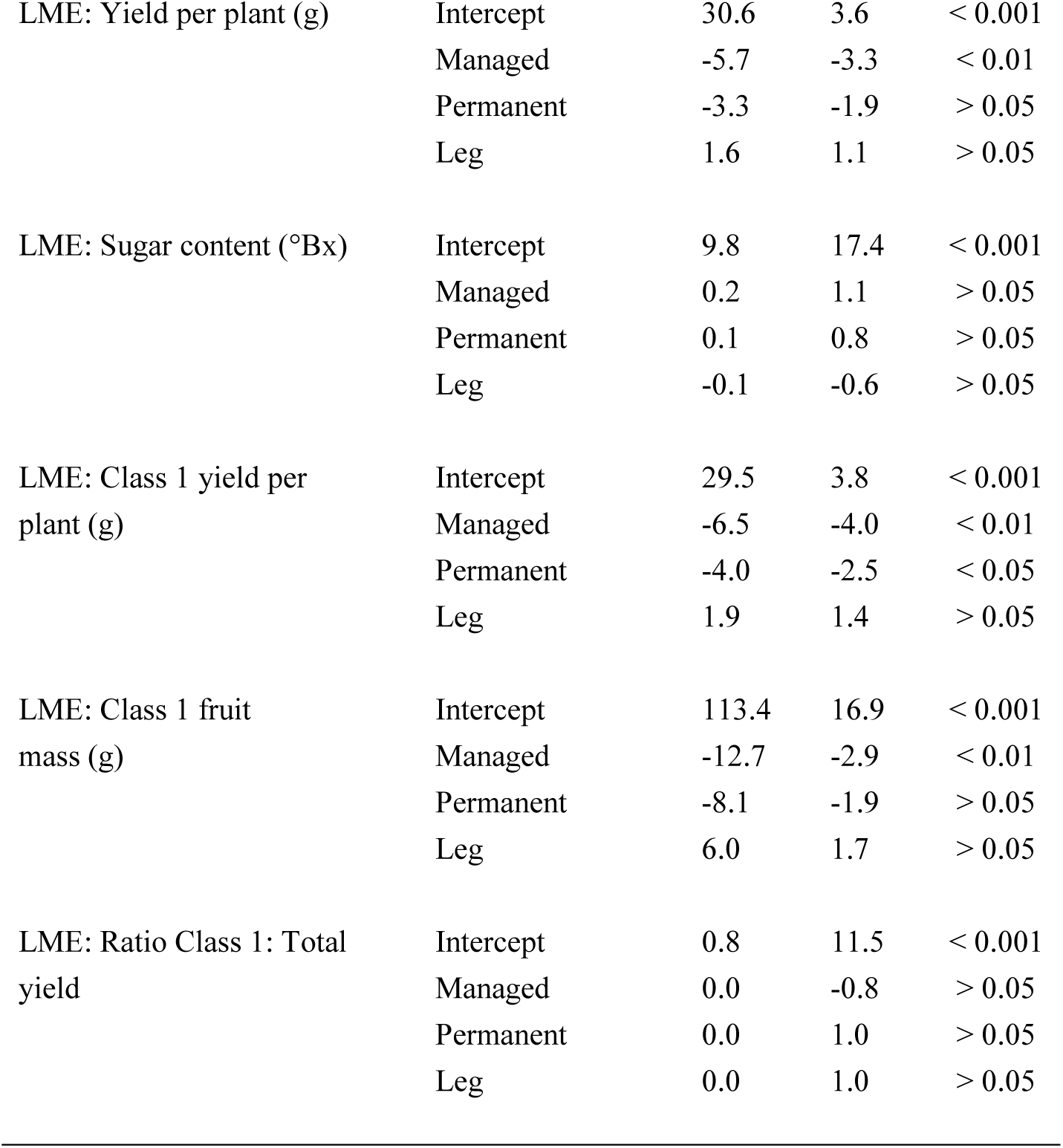
Generalised Linear Model (GLM) outputs for polytunnel environmental conditions; air temperature, relative humidity (RH), photosynthetically active radiation (PAR) irradiance and soil moisture. Explanatory variables are compared with the intercept (control), row, fleece treatment, fleece on/off, day and time of day. Categorical variables are row and fleece treatment. Continuous variables, day and time, are untransformed linear variables unless otherwise stated (cos = cosine, sin = sine). The error structure is gaussian. Linear Mixed Effects (LME) model outputs for treatments effects on fruit yield and quality. Fixed effects were the explanatory variables; fleece treatment (managed and permanent) and row (leg or centre). The random effect was harvesting date to account for the repeated measures design.

Although the high level of replication resulted in low P values across all variables and responses, the relative effect sizes (model values) indicated the degree to which the stated parameter influenced polytunnel conditions. Air temperature and RH increased and decreased over the experimental period, respectively, but the relatively low effect sizes for PAR irradiance and soil moisture, when compared with their absolute values, indicated that mean daily values remained relatively consistent over the period. The fleece treatment was associated with an increase in air temperatures by a mean of 0.8°C and 0.7°C and a decline of 1.4% and 1.8% for RH for the managed and permanent fleece, respectively, across the whole experiment. Direct comparison of only the occasions where the fleeces were on for the different treatments revealed that the air temperature was a mean of 14% higher under the fleeces compared to the control (y = 1.14x, r^2^ = 0.96) and RH was a mean of 7% lower compared to the control (y = 1.07x – 5.9, r^2^ = 0.98). The effect size for the fleece treatments on PAR irradiance were low when compared to their absolute values. Fleecing was associated with increased soil moisture, however, the degree of fit for this parameter was much lower than for air temperatures and RH. Direct comparison of only the occasions where the fleeces were on for the different treatments revealed that soil moisture was a mean of 8% higher than the control (y = 1.08x, r^2^ = 0.77). Although row was significant across the GLM analyses, the effect size for this variable was low across all four of the environmental variables.

### 3.2. Coupling weather station measurements to polytunnel conditions

Trigonometric functions best explained variation in air temperature during the day for the weather station (r^2^ = 0.38, P < 0.001) and polytunnel (r^2^ = 0.63, P < 0.001, Figure 2a). The fitted lines revealed that mean overnight temperatures at the weather station were moderately higher than the polytunnel, however, the polytunnel warmed more rapidly than the weather station and reached a substantially higher maximum temperature at approximately 14:00-15:00. This variation meant that there were differences between air temperatures measured at the weather station and in the polytunnel, with the fitted linear regression explaining 63% of the variation in the dataset (Figure 2b). Trigonometric functions also optimally described the cyclical air temperature differential (polytunnel temperatures – weather station temperatures, r^2^ = 0.62, P < 0.001, Figure 2c). Combining the linear regression equation, which predicts polytunnel air temperature from weather station air temperature, with the equation for the air temperature differential created a simulated polytunnel air temperature (Figure 2d). This simulation explained substantially more variation in actual polytunnel measurements than the untransformed weather station data (r^2^ = 0.83, P < 0.001).

**Figure 2:**
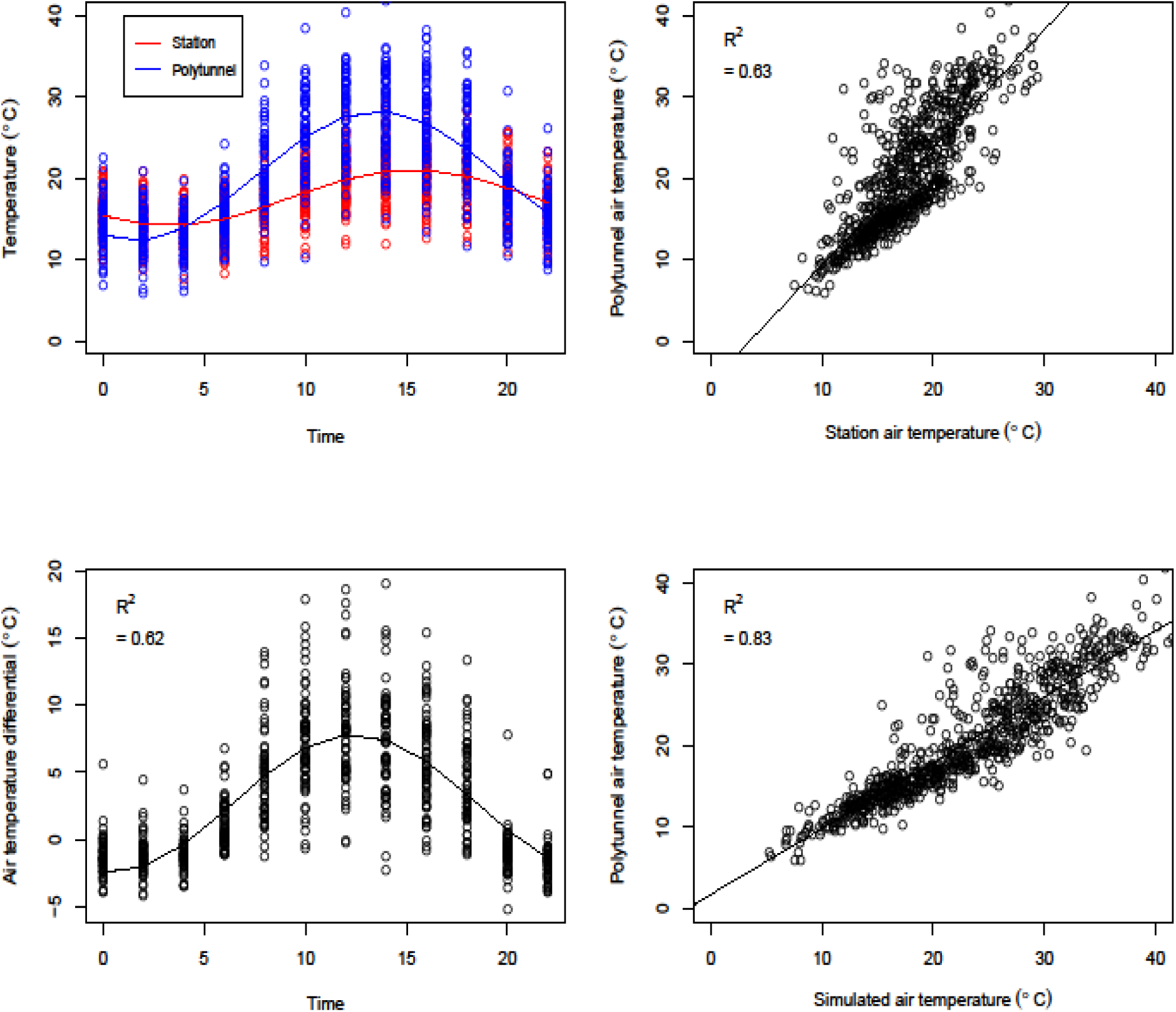
Plots of (a) daily air temperature measurements in the polytunnel (mean of six control sensors) and weather station air temperatures averaged every two hours, (b) weather station and polytunnel air temperatures, (c) daily air temperature differentials (polytunnel temperatures – weather station temperatures) averaged every two hours and (d) simulated air temperatures and measured polytunnel air temperatures (Simulated air temperature = ((1.4*Weather Station)-5.1)+((−0.6*sin(2 * pi/24 * Time)) + (−5.1*cos(2 * pi/24 * Time)) + 2.7, N = 760))

Trigonometric functions also best explained variation in RH during the day for the weather station (r^2^ = 0.29, P < 0.001) and polytunnel (r^2^ = 0.71, P < 0.001, Figure 3a). RH at the weather station was generally higher overnight than the polytunnel but lower during the day, again following a cyclical pattern. The degree of scatter in the dataset meant that there were greater differences between RH measured at the weather station and in the polytunnel than for air temperatures, with the fitted linear regression explaining 52% of the variation in the dataset (Figure 3b). Trigonometric functions also optimally described the cyclical RH differential, with the fitted line explaining slightly more variation that for air temperature (r^2^ = 0.64, P < 0.001, Figure 3c). When the linear regression equation and differential equation was combined the simulated polytunnel conditions explained substantially more variation in actual polytunnel RH measurements than the untransformed weather station data (Figure 3d, r^2^ = 0.83, P < 0.001).

**Figure 3:**
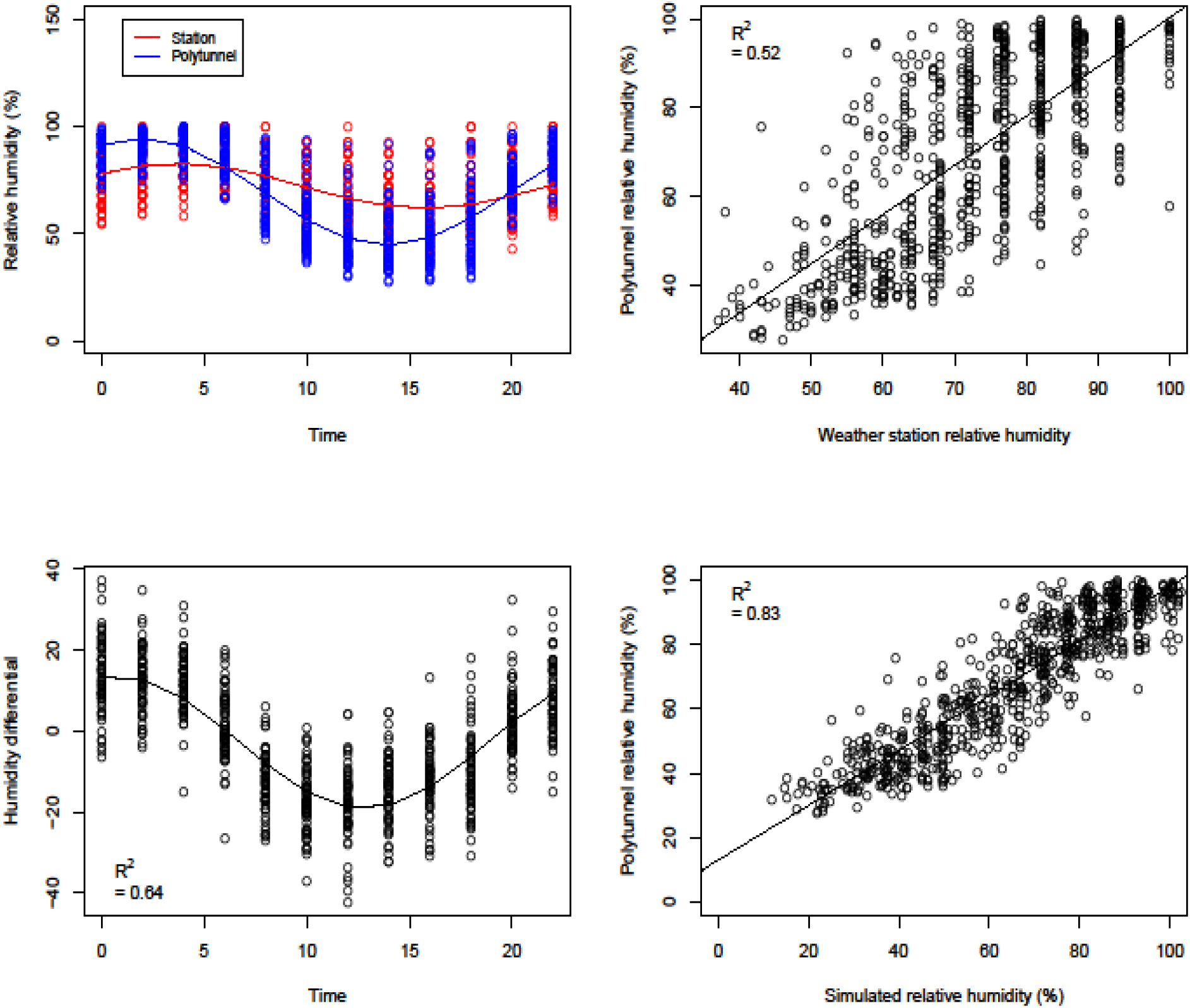
Plots of (a) daily relative humidity measurements in the polytunnel (mean of six control sensors) and weather station relative humidity averaged every two hours, (b) weather station and polytunnel relative humidity, (c) daily relative humidity differentials (polytunnel RH – weather station RH) averaged every two hours and (d) simulated relative humidity and measured polytunnel relative humidity (Simulated RH = ((1.1*Weather station)-10.9)+((3.4*sin(2 * pi/24 * Time)) + (16.0*cos(2 * pi/24 * Time)) −2.8, N = 760))

### 3.3. Defining plant phenology, physiology, fruit yields and quality

The managed fleece treatment was associated with a decline in fruit yield per plant and both fleece treatments were associated with a decline in class 1 fruit yield per plant (Table 1). The managed fleece was also associated with a decline in mean class 1 fruit mass. The permanent fleece treatment was not associated with a difference in yield per plant or class 1 fruit mass when compared to controls, however, P values were less than 0.07 in both cases. Analysis of each of the 15 harvest dates separately revealed that significant declines in fruit yield per plant were detected between the control and both fleece treatments on days 72, 75 and 79, and for the managed fleece on day 83 since planting (all P < 0.05). On these occasions the mean decline in fruit yields per plant were 26.2g and 18.9g for the managed and permanent fleece, respectively. There was one occasion where the fleece treatments produce higher yields than the control, but this only occurred on day 94 for the managed fleece and on day 101 for the permanent fleece, with yield increasing by approximately 2g for both fleece treatments. No other significant differences were detected for the rest of the harvest dates. There were no differences detected between central or leg rows for any of the fruit mass or quality metrics (all P > 0.05).

Stomatal conductance values increased when the fleece was on, by a mean of 130 mmol m^−2^ s^−1^ (t = 5.0, P < 0.001) but there was no effect of row (P > 0.05). LME analysis revealed that stomatal conductance increased by 31 mmol m^−2^ s^−1^ for a 1 °C increase in air temperatures (t = 31.0, P < 0.001) and by 2.4 mmol m^−2^ s^−1^ for a 1% increase in soil moisture (t = 2.1, P < 0.05), but reduced by 14.7 mmol m^−2^ s^−1^ for every 1 °C increase in soil temperatures (t = −3.0, P < 0.01). RH and PAR irradiance were not directly linked to rates of stomatal conductance (P > 0.05).

Logistic growth curves were used to describe cumulative fruit yield per plant over time for the managed fleece (Y_0_ = 0.0026, µ = 0.17, K = 408, r^2^ = 0.87, P < 0.001, figure 4a), control (Y_0_ = 0.0008, µ = 0.19, K = 507, r^2^ = 0.91, P < 0.001, figure 4b) and permanent fleece (Y_0_ = 0.0033, µ = 0.17, K = 452, r^2^ = 0.97, P < 0.001, figure 4c) treatments. High r^2^ values indicated a high degree of fit for cumulative fruit yields over time using logistic growth curves. Similarly, cumulative class 1 fruit yields also followed a logistic growth curve for the control, managed fleece and permanent fleece. Fruit sugar content increased over time (y = 0.2x – 2.8, r^2^ = 0.61, P < 0.001), however, there were no treatment effects in this case (Figure 4d).

**Figure 4:**
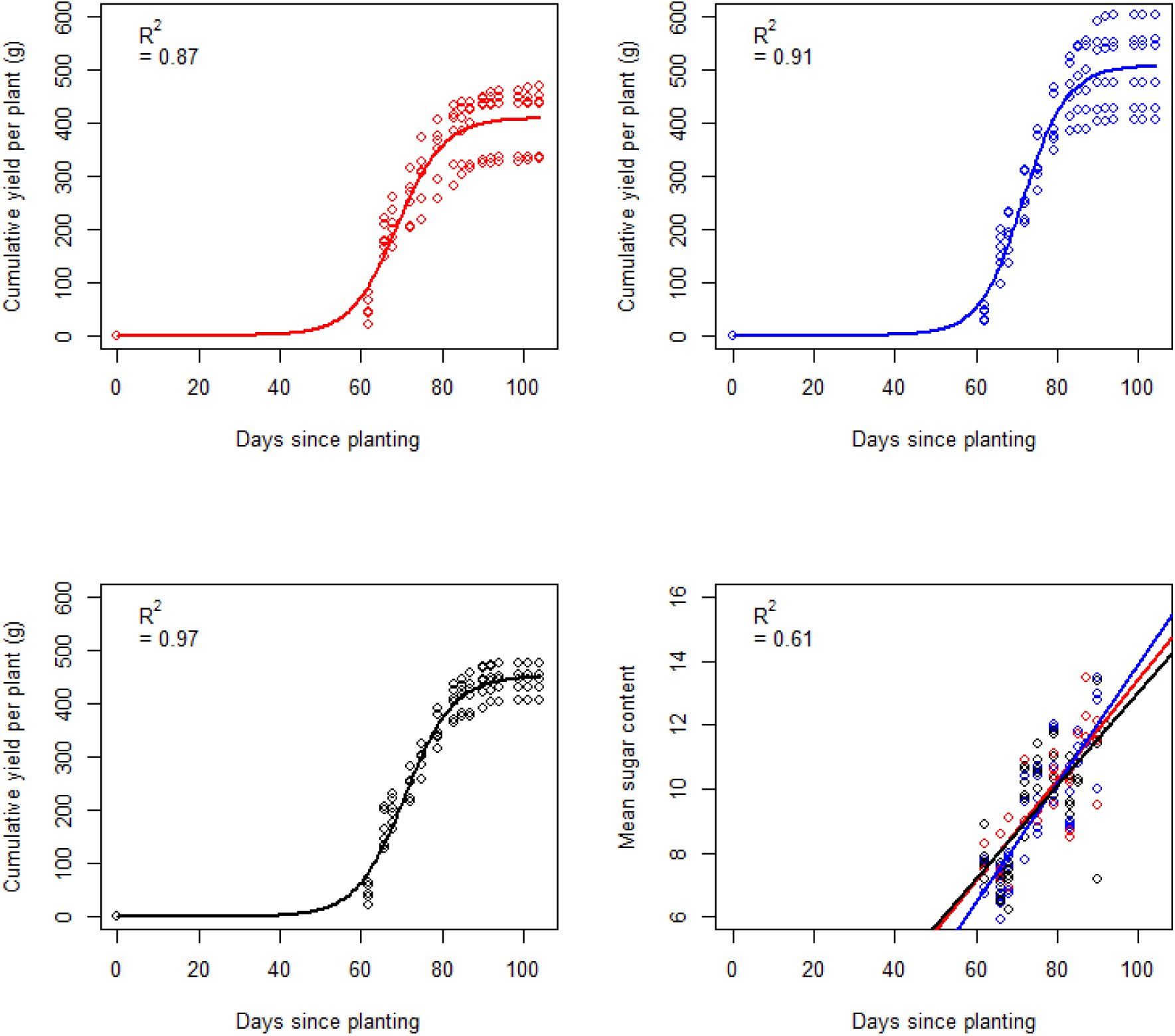
Plots of cumulative fruit yields over time with fitted logistic curves for (a) managed fleece (Y_0_ = 0.0026, µ = 0.17, K = 408), (b) control (Y_0_ = 0.0008, µ = 0.19, K = 507) and (c) permanent fleece (Y_0_ = 0.0033, µ = 0.17, K = 452), and a plot of (d) mean fruit sugar content over time with fitted linear lines (y = 0.2x – 2.8). Each point represents mean fruit yield or sugar content for one of six dataloggers for each treatment and for each harvest (N = 90)

### 3.4. Coupling phenology model and datalogger observations using machine learning

Phenology model parameters (r, K and d) were optimised using machine learning by gradient descent and fit estimated with least-squares. Adequate convergence of parameter estimates was observed after 10,000 steps across each of 10 replicates (repeat machine learning runs). The fitted phenology model parameters for each experimental block (N = 17 blocks after removing one block due to missing data) were coupled to the datalogger observations (PAR, RH and mean air temperature) using GLMs to predict each growth model parameter in turn (Equations 2, 3 and 4). For each, a maximal model with interaction of the form (<parameter> ~ mean_temperature * RH * PAR) was considered initially. Thereafter, insignificant terms and interactions (F-ratios) were refined using reverse stepwise (sequential) deletion via AIC to produce a minimal model for each parameter. Fitted models were checked for homoscedasticity and normality of variance. In each case RH proved to be the most significant parameter and the inclusion of RH alone into the fitted logistic growth equations was the optimal fit for the phenology measurements (Table 2).

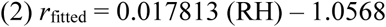

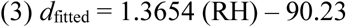

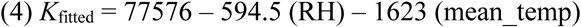

**Table 2:** Minimum fitted phenology model parameters when coupled to the datalogger observations (PAR, RH and mean air temperature) using General Linear Models (GLMs) to predict each growth model parameter in turn. Phenology model parameters (r, d and K) were optimised using machine learning by gradient descent and fit estimated with least-squares. Convergence of parameter estimates was observed after 10,000 steps.

| Parameter | Estimates | Pr ($>\vert t\vert $) | F (df) |
| --- | --- | --- | --- |
| $r$ | Intercept: -1.0568 | $< 0.05$ | 7.71 (1,15) |
| | Coefficient, RH: 0.0178 | $< 0.05$ | |
| $d$ | Intercept: -90.23 | $< 0.001$ | 10.33 (1,15) |
| | Coefficient, RH: 1.3654 | $< 0.001$ | |
| $K$ | Intercept: 77576 | $< 0.05$ | 3.95 (2,14) |
| | Coefficient, Temperature: -1623 | $< 0.05$ | |
| | Coefficient, RH: -594.5 | $< 0.05$ | |

Equations (2), (3) and (4) may be substituted into equation (1) as parametisations for r, d, and K in terms of RH. In this sense they are effectively link functions relating the climatic conditions to the growth model. For this study only RH was needed for a sufficient model, but in other contexts the coefficients and explanatory variables present in the link functions would change – but not the key growth model in equation (1).

### 3.5. Agronomic implications

This study presented and tested a novel framework for linking mesoscale meteorological forecasts to local crop microclimates using embedded autonomous sensors and machine learning. This approach transformed mesoscale meteorological forecasts into site- and polytunnel-specific predictions for temperature and RH. There was natural variation in growing conditions within the polytunnel, but addition variation was generated using fleece applications. We have demonstrated that fruit phenology follows logistic growth curves over time and the coefficients of these curves are dependent on polytunnel conditions, namely RH in this case. We have shown that machine-learning techniques can be used to calculate these coefficients and in doing so parametrise models which can predict fruit yields and timing to a greater degree of accuracy that previously possible. This has the potential to solve a key agronomic challenge because predicting yields, quality and phenology is important for many soft-fruit growers, since fruits are perishable, high-value and seasonal, and harvesting is labour-intensive and reliant on an increasingly expensive workforce (Morris et al. 2017).

The features and relationships in the framework follow from simple and tractable agronomic and meteorological principles that can, in principle, be used for predictions of phenology by coupling the mesoscale forecast directly to the phenology model without the use of dataloggers to localise abiotic data. However, in practice these models’ coefficients will vary according to many factors, including weather station proximity, polytunnel design, slope, elevation and orientation, and other site characteristics. This framework would therefore use iterations over several crops in the year, and over a number of years, to train model parameters to represent geographic, seasonal and inter-annual variation. Each model will necessarily be bespoke for an individual site, crop and agricultural system but can be generated using a consistent framework. In this case, the weather station was 3 km from the site and the orientation of the polytunnel was north to south. The physical design of a polytunnel and constant geographical relationship to the sun allows for cyclical residual differences between forecasted and actual growing conditions which can be modelling using trigonometric functions. Yields and phenology responses to growing conditions will, of course, vary between crop types and varieties (Grant et al. 2012; Vicente et al. 2014) but our framework allows for the development of predictive models without *a-priori* crop phenology knowledge.

Dataloggers can be installed to a growing location at the start of the growing season or over the winter and used to collect localisation data (e.g. microclimate adjustment parameters) in conjunction with the mesoscale observations available from a weather station. Datalogger observations can also be used in combination with yield data after the first few crop cycles (~65d each) to infer both the phenology model parameters (for a single point in the growing season) and seasonality adjustment factors to the basic phenology model (e.g. adjustments arising from the time of year and climatic variation). After sufficient training the end-to-end phenology model system will be adequately trained to predict yields and timings based only on the mesoscale weather forecast data, though dataloggers could optionally be retained to permit real-time adjustments to the phenology model predictions. Farm managers could intervene to slow down to speed up ripening depending on the terms of their agreements with their customers and staffing. Extensions to the modelling procedure here could use decision forest learning algorithms or neural networks to couple phenology model, datalogger, and localisation model parameters, instead of multivariate parametric model selection and fitting (Ghahramani 2015).

Our framework could be readily generalised to other crops and climates and further parametrised with real-time imaging data. This also demonstrates the applicability of machine-learning and/or numerical optimisation approaches to agriculture. Wide adoption of autonomous sensor technology in the future will yield secondary science benefits, generating rich time-series data to mine for correlates of climate change, leading to enhanced predictions of climate change susceptibility to inform climate resilience and adaptation measures. These datasets can also feed back into an enhanced understanding of modified varieties’ real-world performance in relation to microclimates, mycorrhizal fungi, pests and diseases (Camprubi et al. 2007; Xu et al. 2012). Powdery mildew infection risk, for example, is likely to be predictable using environmental data (Carisse et al. 2013). In the case of this strawberry variety, Malling Centenary, RH was the strongest predictor of field yields and phenology. This is contrary to the common use of air temperature in other fruit phenology models, however, such models have been shown to be imprecise – for example, GDD has been shown to range by 297–347 within varieties (Krüger et al. 2012). It should be noted that RH is a function of air temperature (warmer air can hold more water) and increased RH has been correlated with increased fruit firmness (Pyrotis et al. 2012), fruit size (Lieten 2002) and nutritive values (Shin et al. 2007). Relative humidity may be a useful metric for predicting fruit phenology and yields because it is an integrative metric of water status and temperature. Increased knowledge of varietal responses to growing conditions will not only assist with farm management and profitability but could also contribute to the selection and breeding of optimal varieties to cultivate under global environmental changes (Kimball et al. 2007; Luedeling et al. 2011).

Our experimental fleece treatments were introduced to generate variation in fruit yields and phenology for machine learning purposes. However, the extent of its use was not realistic, since fleeces are currently used transiently to prevent frost damage on commercial strawberry farms. Some of our findings, however, are of value to agronomists and growers since we discovered that applying the fleece generally increased air temperatures by 14%, reduced RH by 7% and increased soil moisture by 8% and made small differences to PAR. PAR was not a strong predictor of phenology in this study, but raised PAR has been shown to reduce the period from anthesis to fruit harvest elsewhere, and is likely to become incorporated into predictive models when there is a greater degree of spatial variability in PAR values (Krüger et al. 2012). Increased air temperatures may have been caused by the insulator effect of the fleece material and increased soil moisture may be driven by reduced evaporation from the soil surface. RH declined with rising air temperatures, but by a lesser extent, indicating that additional water may be released by the plants, as shown by the stomatal conductance measurements. Fleecing reduced total yield and class 1 fruit yield, and increased rates of stomatal conductance. This was linked to greater air temperatures and soil moisture content and is indicative of heat stress (Martínez-Ferri et al. 2016). Embedded autonomous sensors and coupled weather models may therefore also be used by farm managers to prevent acute problems caused by growing conditions, such as adding fleeces only during the periods when frost is likely or opening doors and vents to protect plants if severely high temperatures are anticipated.

### 3.6. Limitations

A current challenge in the automation of our machine learning framework is our reliance upon manually gathered fruit yield and quality data, as well as manual physiological measurements. We measured changes to plant physiology, represented by stomatal conductance, in response to changes in growing conditions and application of the fleece treatments. Changes to plant physiology may be the underlying mechanism for changes in fruit phenology and yield that we observed, as has been proposed elsewhere (Grant et al. 2012). However, there is currently no cost-effective way of reliably measuring this in real-time across a sufficiently large number of individuals. There is emergent technologies, including thermal imaging (Grant et al. 2016), high resolution cameras (Delalieux et al. 2017) and wearable sensors made of graphene based nanomaterials (Oren et al. 2017) which may provide us with the resolution of data that is necessary to provide a greater mechanistic understanding of changes to yields and phenology. If these technologies are coupled with our machine learning framework then our iterative framework of model training could be entirely automated. Nevertheless, at this stage in the development of integrative artificial intelligence and networked microsensors, our approach offers a step forward in coupling weather forecasting to local polytunnel growing conditions and in generating bespoke phenological prediction models to assist fruit growers.

## 4. Conclusion

We have developed a novel framework for linking mesoscale meteorological forecasts to local crop microclimates using embedded autonomous sensors. This parametrises a linked ripening model and predicts crop yields and ripening dates. Our approach transformed mesoscale 2-hourly meteorological forecasts, which showed a relatively low agreement with polytunnel air temperature (R^2^ = 0.6) and RH (R^2^ = 0.5) measurements, into site- and polytunnel-specific predictions for temperature and RH (both simulated R^2^ = 0.8). We have demonstrated that fruit phenology (yields and fruit timing) follow logistic growth curves and the coefficients of these curves are dependent on polytunnel conditions. We present a framework for using machine-learning techniques to calculate these coefficients using embedded autonomous sensors and in doing so parametrise models which can predict fruit yields and timing to a greater degree of accuracy that previously possible. This has the potential to solve a key agronomic challenge because predicting yields, quality and phenology is important for many soft-fruit growers, since fruits are perishable, high-value and seasonal, and harvesting is labour-intensive and reliant on an increasingly expensive workforce.

## Acknowledgements

Thanks to Berry Gardens Ltd for providing polytunnel space, plants and technical support (www.berrygardens.co.uk). We also thank Mothive Ltd for providing data loggers and sensors, installation, data acquisition and data storage (www.mothive.com).

